# Classification of Pacemaker Dynamics in the Mouse Intestine by Field Potential Microimaging

**DOI:** 10.1101/2021.05.29.446256

**Authors:** Naoko Iwata, Chiho Takai, Naoto Mochizuki, Mariko Yamauchi, Noriyuki Kaji, Yoshiyuki Kasahara, Shinsuke Nakayama

## Abstract

The flexible and sophisticated movement of the gastrointestinal (GI) tract implies the involvement of mechanisms other than enteric neural circuits, to coordinate excitation in microregions. We thus performed microimaging of pacemaker dynamics in the small intestine of mice since it contains typical network-forming pacemaker cells. A dialysis membrane-reinforced low-impedance microelectrode array (MEA) enabled field potentials over a wide frequency range to be stably measured in microregions. The pacemaker dynamics were classified into basic patterns despite large variations. In the developmental process, pacemaker activity was categorized as either an ‘expanding’ or a ‘migrating’ pattern that was initiated in or propagated to the MEA sensing area, respectively. The intercellular current of the volume conductor complicated the waveform of both activities. The existence of ‘expanding’ and ‘migrating’ patterns was attributable to duplicated pacemaker systems such as intracellular Ca^2+^ oscillation-activated and voltage-gated mechanisms. Additionally, from the spatio-temporal feature during the period of pacemaker events, the ‘bumpy/aberrant’ pattern was defined by aberrant, incoherent propagation, and associated with local impairment of excitability, while the ‘colliding/converging’ pattern involved the interaction of multiple activities in the MEA area. Interconversion between the four micro-coordination patterns occurred in the same microregion. 5-Hydroxytryptamine (5-HT) promoted ‘migrating’ activity, implying an improvement or restoration of spatial conductivity. These results agree well with the action of 5-HT to change GI movement toward propulsion. In conclusion, our MEA method of microimaging classification enables the quantitative assessment of spatio-temporal electric coordination underlying GI motility, suggesting its application to small model animals.

## 1. Introduction

The flexible and sophisticated movement of the gastrointestinal (GI) tract implies an underlying mechanism that has various forms coordinating spatio-temporal excitability in microregions (Spencer et al. 1999; Morishita et al. 2017), and is simultaneously operated together with the reflex of intrinsic neural circuits (Bayliss & Starling, 1899). The patterns of GI movement, especially in the small intestine, are often described using a few limited terms such as ‘segmentation’ and ‘peristalsis’ for agitation and transportation of the luminal contents, respectively, but in reality, the GI tract exhibits smooth, elaborate and highly varied movements that show subtle changes between cycles, for example as demonstrated using abdominal cine magnetic resonance imaging (Teramoto et al. 2012).

Electrical excitation underlies numerous GI movements (Tomita, 1981; Szurszewski, 1987). In the vast majority of the tract, spontaneous electrical activity originates from special interstitial cells that act as pacemaker cells, which are now recognized for their additional crucial role in coordinating local activities through their network-like structure (Faussone-Pellegrini, 2005; Nakayama et al. 2006; Lammers et al. 2008; Huizinga & Lammers, 2009). However, the progress of spatio-temporal investigation of pacemaker potentials has been slowed by a debate on the waveforms between intracellular and extracellular recordings (Sanders et al. 2016; Huizinga, 2017; O’Grady et al. 2017).

Reflecting spontaneous pacemaker activity, intracellular microelectrodes exhibit relatively simple wave forms comprised of initial rapid depolarizing potential followed by a plateau phase of variable amplitude and duration (Sanders et al. 2016), but simultaneous intracellular recording with multiple microelectrodes requires a masterful handling. On the other hand, extracellular recordings exhibit a wide variety of waveforms such as biphasic and triphasic potentials. In this debate, it has been pointed out that the waveform of extracellular field potential is theoretically proportional to the first-time derivative of the transmembrane voltage, but in practical measurements, this relationship seems to be more complicated (Sanders et al. 2016; Huizinga, 2017; O’Grady et al. 2017). It has also been discussed that the complex waveforms in extracellular recording may originate from recording and data processing techniques, including electric filters, and/or variations of active excitable cells, depending on experimental conditions such as use of L-type Ca^2+^ channel blockers that abolish transient action potentials from smooth muscle cells (Dickens et al. 1999; Akbarali et al. 2010).

In this study, we thus performed microimaging of pacemaker field potentials in order to interpret the complex waveforms of extracellular recordings. We employed a low-impedance microelectrode array (MEA) fabricated with nanoparticles, and firmly mounted samples with dialysis membranes. This method enabled field potentials over a wide range of frequencies to be stably measured in microregions. In other words, this method solved the electric discrepancy underlying the decrease in efficacy of electric signal transmission as electrode size decreases, and repeatedly monitored pacemaker dynamics in the same microregion.

In the small intestine of mice, despite huge variations in each pattern, field potential images were classified into two major micro-coordination patterns, ‘expanding’ and ‘migrating’, depending on the developmental process whether pacemaker activity was initiated in or migrated to the MEA area. In both ‘migrating’ and ‘expanding’ activities, a characteristic feature is that positive potentials precede negative potentials during the propagation in the MEA area, providing a rationale for the complicated waveforms of extracellular recordings in the GI tract. Due to its syncytal nature (Tomita, 1970; Dickens et al. 1999; Manchanda et al. 2019), an intercellular pacemaker current also participates in the depolarization of the membrane during an early phase of propagation, and flows in the opposite direction against the transmembrane pacemaker current. In addition, 5-hydroxytryptamine (5-HT, serotonin), which is known to promote propulsive movements (Bülbring and Lin, 1958; Mawe and Hoffman, 2013), exerted consistent modulatory effects on micro-coordination patterns *in vivo*, suggesting that this MEA technique is beneficial for functional assessment of spatio-temporal electric activity.

## 2. Materials and Methods

### 2.1. Animals and preparations

Animals were treated ethically in accordance with the guidelines for the proper conduct of animal experiments published by the Science Council of Japan. All procedures were approved by the Animal Care and Use Committee of Nagoya University Graduate School of Medicine (permissions #28316, #29121, #30312 and #31199). C57BL/6J wild-type mice (8–40 weeks, either sex) and *W/W*^*v*^ (WBB6F1/Kit-*Kit*^*W*^*/Kit*^*W-v*^/Slc) mice (8–12 weeks, male, Japan SLC, Hamamatsu, Japan), which lack interstitial cells of Cajal (ICCs) in the myenteric plexus (MP) region of the small intestine, were sacrificed by cervical dislocation after the inhalation of carbon dioxide. The whole GI tract was quickly resected, and small pieces (~20 mm) of the ileum were isolated and cut along the mesentery. The entire muscularis propria containing the myenteric plexus was isolated by removing it from the mucous membrane using forceps.

### 2.2. Immunohistochemistry

Small segments of the isolated ileal musculature were fixed with ice-cold acetone for 10 min, then blocked with 2% bovine serum albumin for 1 h. Samples were incubated with anti-c-kit rat monoclonal antibody (1:500 v/v, ACK2, BioLegend, San Diego, CA, USA) overnight (4ºC), and subsequently stained with donkey anti-rat IgG antibody, with Alexa Fluor 488 conjugate (1:500 v/v, A-21208, Thermo Fisher Scientific, Waltham, MA, USA). Fluorescence images were acquired from 70 μm thick slices under a confocal microscope (TCS SP5 II, Leica Microsystems, Tokyo, Japan) using a ×20 objective lens.

### 2.3. Solutions and drugs

The extracellular solution used for the MEA recordings was a modified Krebs solution with the following composition (in mM): NaCl 125, KCl 5.9, MgCl_2_ 1.2, CaCl_2_ 2.4, glucose 11, and Tris-HEPES 11.8 (pH 7.4). Nifedipine and 5-HT hydrochloride were purchased from Sigma-Aldrich (St. Louis, MO, USA).

### 2.4. Electrical recordings

We developed the dialysis membrane-reinforced MEA recording technique to stably monitor the micro-organization of ICC pacemaker activity in the GI tract of small animals (Iwata et al. 2017; Morishita et al. 2017) because many animal disease models are developed using small animals. In addition, there is no consensus for the application of multi-electrode arrays used previously to small animals (Lammers et al. 2008; O’Grady, 2012; Sanders et al. 2016; Huizinga, 2017; O’Grady et al. 2017). To demonstrate the efficiency of MEA field potential imaging, pacemaker potentials were first measured in the presence of nifedipine (2 μM), which selectively blocks L-type voltage-gated Ca^2+^ channels and suppresses smooth muscle contraction (Dickens et al. 1999; Akbarali et al. 2010). Additionally, the effect of 5-HT was examined as a proof of concept that this method can be applied for functional assessment of spatio-temporal electric activity in the GI tract.

An 8 × 8 array of microelectrodes (P515A, Alpha MED Scientific, Ibaraki, Japan) was used to measure electric activity in the ileum. The inter-electrode distance of the MEA was 150 μm, and the sensing area was 1050 × 1050 μm^2^. Samples of mouse ileal muscle sheets were firmly mounted onto the MEA, with the longitudinal muscle layer facing downward, using a small piece of dialysis membrane (cellulose tube, Visking, Chicago, IL, USA) and a slice anchor (SDH series, Harvard Apparatus Japan, Tokyo, Japan). A large reference electrode made of platinum was immersed in the extracellular solution (Fig. 1A, B). During the propagation of pacemaker potentials, the membrane currents (*I*_M_) flowing through the access resistance (*R*_a_) between the sensing and reference electrodes changed the field potential (*I*_M_ × *R*_a_) (Fig. 1B). A set of 8 × 8 field potentials (Fig. 1C, E) originating from c-Kit–positive interstitial cells (i.e. ICC-MP, Fig. 1D) was simultaneously recorded using a multi-channel AC amplifier that stabilized the baseline potential with high-pass filtering at 0.1 Hz and reduced noise with low-pass filtering at 10 kHz (Fig. 1E) in accordance with conventional extracellular electrode recording techniques (Astrand et al. 1988). The arrayed data were digitized and stored on a personal computer using a 14-bit A/D converter with a sampling rate of 20 kHz (50-μs intervals). The dynamic range of A/D conversion was set to ± 1 mV with a digital resolution of ~0.12 μV.

**Fig. 1.**
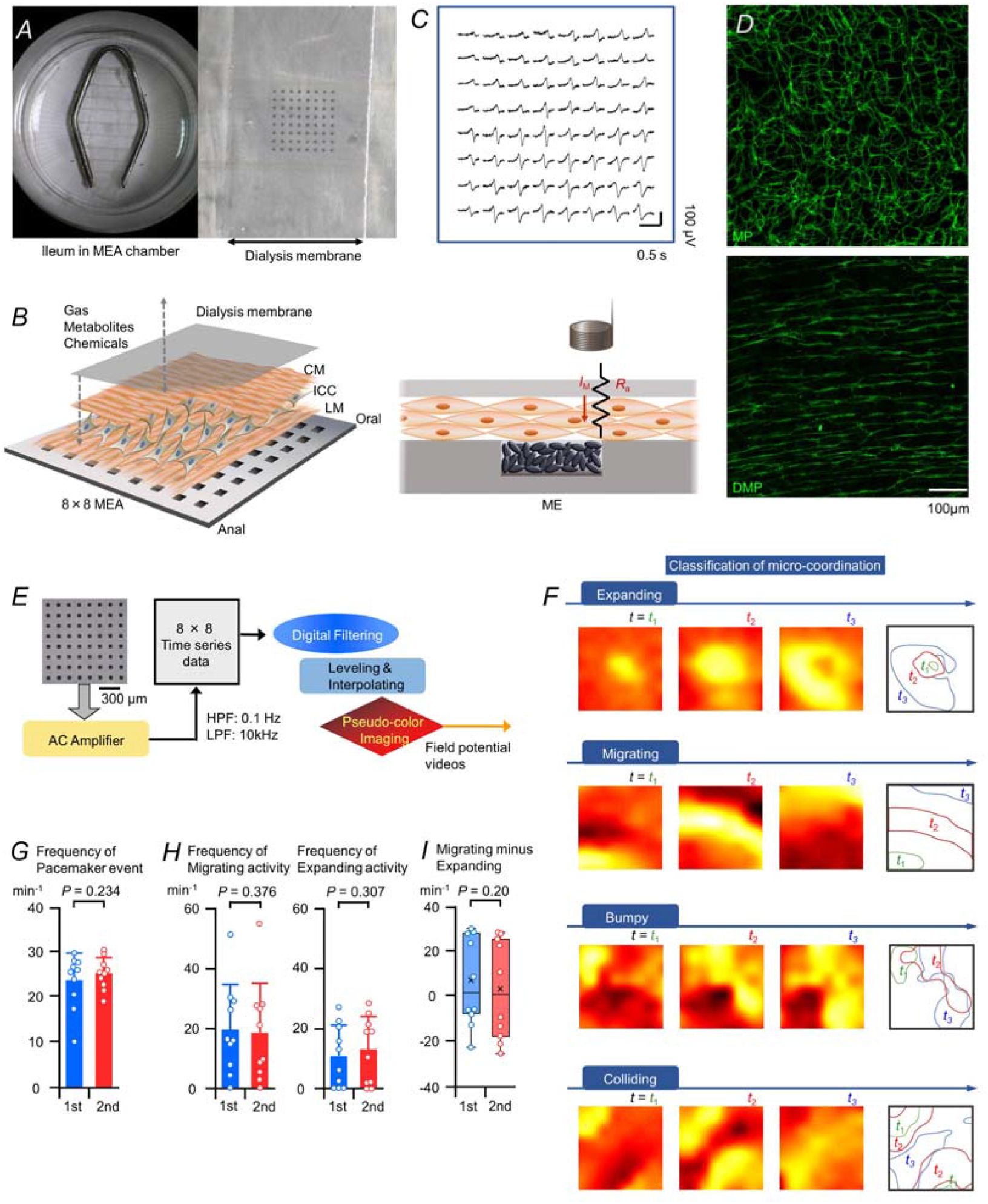
Dialysis membrane-reinforced microelectrode array (MEA) recordings enabled classification of pacemaker micro-coordination. (**A**) Photographs showing a muscle sample isolated from the mouse ileum and mounted on an 8 × 8 MEA using a piece of dialysis membrane and an anchor. (**B**) Illustration showing the arrangement of the muscle sample and dialysis membrane on the MEA plate. The muscle sheet was positioned with the longitudinal muscle layer facing downward and with the oral and anal ends located toward the upper and lower sensing electrodes, respectively. Each microelectrode (ME) sensed changes in field potential reflecting the flow of local membrane current (*I*_M_) through the access resistance (*R*_a_) between the ME and a reference electrode. CM: circular muscle; ICC: interstitial cells of Cajal; LM: longitudinal muscle. (**C**) An 8 × 8 plot showing the spontaneous electric activities recorded from a muscle sample in the presence of nifedipine. (**D**) Confocal images of c-kit immunofluorescence showing ICCs in the myenteric plexus region (MP: upper) and the deep muscular plexus region (DMP: lower) (representative of ≥3 experiments). (**E**) Procedures used for potential mapping. HPF: high-pass filtering; LPF: low-pass filtering. (**F**) Patterns of pacemaker potential micro-coordination: ‘expanding’, ‘migrating’, ‘bumpy/aberrant’ and ‘colliding/converging’ (see text and Supplemental Fig. S1 for definitions). (**G, H**) Graphs showing the frequency of pacemaker events (min^-1^) and frequency of ‘migrating’ or ‘expanding’ activity (min^-1^) during the first and second halves of the MEA measurements (54–90 s) in 10 samples. (**I**) Box-and-whisker plot of the difference between the frequency of migrating and expanding activity (min^-1^) in **H**. ×, the mean; vertical line, the median value.

For all samples, the oral and anal ends of the muscles were oriented toward the upper and lower ends of the MEA, respectively. The dialysis membrane had a molecular weight cut-off of 12,000, allowing oxygen, ions and energy metabolites to be exchanged. The MEA recording chamber (~2 mL volume) was placed on a heater maintained at ~34ºC. The samples were incubated at least 20 min, and MEA recording was started after the occurrence (frequency) of pacemaker activity reached a quasi-stable state. Nifedipine (2 μM), a dihydropyridine Ca^2+^ channel antagonist, was added to the extracellular medium to selectively block L-type voltage-gated Ca^2+^ channels, which act as a major Ca^2+^ influx pathway in GI smooth muscle (Akbarali et al. 2010) and suppress smooth muscle contraction without impeding the pacemaker activity of ICCs (Dickens et al. 1999; Kito and Suzuki, 2003). The above procedures stabilized the electric conditions because similar noise levels persisted throughout the measurements, both in the control and in the presence of 5-HT.

Each sensing microelectrode of the MEA (P515A) had a square profile with a side length of ~50 μm and was made from Pt black nanoparticles, which increased the surface area by ~200-fold (equivalent to 0.5 mm^2^) (Fig. 1B). From the capacitance (*C*_ME_: 0.052 μF) and resistance (*R*_ME_: 15 kΩ) of the sensing microelectrode, the estimated impedance was sufficiently low to follow a wide frequency range of electric signals (Taniguchi et al. 2013; Iwata et al. 2017). For instance, impedance (*Z*_ME_) of the sensing electrode at 0.1 Hz [~31 MΩ = √ {1/(2π × 0.1 Hz × 0.052 μF) ^2^ + (15 kΩ)^2^}] was sufficiently small compared with the input resistance of the multi-channel amplifier used in this study (100 MΩ). Thus, the efficacy of electric signal transmission (*Tr*) was estimated to be ~95% at 0.1 Hz [100 MΩ / √ {(100 MΩ)^2^ + (31 MΩ)^2^}].

### 2.5. Field potential data processing

The arrayed field potential data were acquired at a 14-bit depth with a sampling interval of 50 μs. The digital resolution was reduced to 0.5–25 ms to display the time course of pacemaker oscillations. Data processing, including digital filtering and linear spectrum analysis, was performed using commercial software (Kaiseki, Kyowa Electronic Instruments, Tokyo, Japan; Mobius, Alpha Med Scientific, Ibaraki, Japan; MATLAB software package, MathWorks, Natick, MA, USA).

For field potential microimaging (Fig. 1E), after reducing the digital resolution to 5 ms, field potential data were processed by a band-pass filter at 0.25–10.5 Hz. Numerous factors, including variations in *R*_ME_ and *R*_a_ and differences in *I*_M_ between different populations of pacemaker and modulator cells, were considered to affect the magnitude of oscillations in field potential. Therefore, to compensate for these factors and reconstitute a smooth field potential map, the amplitude of the field potential recorded in the n^th^ microelectrode region [ME(n)] was corrected by a normalizing factor [*F*_N_(n)]:

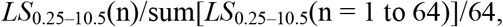

where *LS*_0.25–10.5_(n) is the density of the linear spectrum in the frequency range between 0.25–10.5 Hz for the field potential data at ME(n). To avoid excessive compensation, *F*_N_(n) was limited to within 1.0 ± 0.5. The sum of the *LS*_0.25–10.5_ values for all 64 MEs was used to represent the magnitude of pacemaker activity. Subsequently, the 8 × 8 field potentials were interpolated with 50 points using a ‘cubic’ protocol, and each point of the field potential map was displayed as a ‘hot’ map (MATLAB software package) (Morishita et al. 2017).

### 2.6. Classification of micro-coordination patterns

The series (video) of field potential images (for 54 – 90 s) were used to classify the spatio-temporal features of pacemaker potentials at ‘event’ and ‘activity’ levels (Supplemental Fig. S1; Fig. 1F). In each pacemaker period (i.e., pacemaker event), the micro-coordination of pacemaker potentials was classified at three check points. First, the pacemaker event was assessed to appreciate whether pacemaker potentials propagate in a coherent or incoherent/aberrant manner (Check-1). Second, the event was assessed to verify whether a pacemaker event contains single or multiple pacemaker activities (Check-2). Third, each activity was assessed to observe whether it occurs outside or inside the MEA sensing area (Check-3).

During the pacemaker period, if pacemaker potentials propagated in an aberrant (i.e., extraordinarily incoherent) manner, the event was classified as a ‘bumpy/aberrant’ pattern event. On the other hand, an event showing coherent/regular propagation was classified by the following two check points.

In the first check point, if the event showing coherent/regular propagation contained single independent pacemaker activity, the activity was classified by the region where activity occurs. Consequently, propagation as a volume conductor from adjacent regions outside of the MEA sensing area was defined as ‘migrating’ activity, while activity initiated in the MEA sensing area was defined as ‘expanding’ activity. In addition, ‘migrating’ and ‘expanding’ activities in a single activity event were defined as ‘migrating’ and ‘expanding’ events, respectively.

In the second check point, if the event showing coherent/regular propagation contained multiple pacemaker activities originating from different regions, the event was classified as a colliding/converging pattern. In addition, each activity seen in the event was classified into ‘migrating’ or ‘expanding’ activity, identical to the classification of a single activity event, outside or inside the MEA.

In the ‘bumpy/aberrant’ event, pacemaker activity was classified into a ‘migrating’ or ‘expanding’ pattern activity when the number of activities was countable, i.e., whether the event contained single or multiple activities. This special case was not observed in the series of MEA experiments that examined the effects of 5-HT. Of note, the number of pacemaker event patterns may be increased in a future study. For example, use of machine learning software may reveal complex patterns of spatio-temporal coordination.

In the statics of pacemaker activity pattern, each activity observed in ‘bumpy/aberrant’ and ‘colliding/convergence’ events was counted as a ‘migrating’ or ‘expanding’ activity. For example, in the case that a ‘colliding/converging’ event contained two pacemaker activities showing a ‘migrating’ or ‘expanding’ pattern in the MEA sensing area, 0.5 ‘migrating’ and 0.5 ‘expanding’ activities were counted per event (e.g. Fig. 4G). It is considered that the ratio of ‘migrating’ to ‘expanding’ activity (R_M/E_) can approximately estimate the area covered by each initiating site of pacemaker activity (A_iPA_): MEA sensing area × (R_M/E_ + 1) (mm^2^).

### 2.7. Motion tracking

To assess the mechanical influence on electrical recording, transmission images were acquired by a video camera system in an inverted microscope. After MEA measurement of pacemaker potentials were performed in the presence of nifedipine (2 μM), dialysis membrane-reinforced ileal muscle sheets were placed in a glass-bottomed chamber containing the same extracellular solution. The glass-bottomed chamber was placed on a heating glass (34ºC), and transparent images were acquired. Measurement of transverse (X) and vertical (Y) shifts of a square tracking region from its initial position using motion tracking software (2D-PTV, Digimo, Osaka, Japan) showed negligible movement (Supplemental Fig. S2).

### 2.8. Statistics

Numerical data are expressed as the mean ± S.D. (relative standard deviation: RSD = S.D./|mean|). Comparisons of normally distributed datasets were made to estimate the probability (*P*) of significant differences using paired *t*-tests (*P* < 0.05), except for comparisons of patterns using χ^2^ tests in Table 1. In the control condition, a comparison between the first and second half of each MEA measurement was made in 10 muscle samples from 10 mice (eight males and two females, 8-36 weeks-old mice). The effect of 5-HT was analyzed using MEA measurements that were acquired approximately 5 min after a bath application of 5-HT in nine mice (male, 8-18 weeks-old mice).

**Table 1.** Stability of the occurrence of pacemaker activity between ‘migrating’ and ‘expanding’ patterns in a musculature sample monitored in normal solution for a long duration.

| Time after incubation in MEA chamber | Frequency of migrating activity (min<sup>-1</sup>) | Frequency of expanding activity (min<sup>-1</sup>) | *P* value ($\chi^2$ test) |
| --- | --- | --- | --- |
| 25 min | 9.8936 | 18.829 | 0.71 |
| 40 min | 9.0332 | 20.002 |  |

## 3. Results & Discussion

### 3.1. Patterns of pacemaker micro-coordination

Our dialysis membrane-reinforced low-impedance MEA technique enabled electric activity in the GI tract of small animal models to be stably monitored. Smooth muscle sheets isolated from mouse ileum were mounted on an 8 × 8 MEA with an interpolar distance of 150 μm and a sensing region of ~1 mm^2^. Each smooth muscle sheet was mounted beneath a piece of dialysis membrane that maintained the conditions required for electric recording, such as access resistance (*R*_a_) between the sensing and reference electrodes and isolation resistance between the sensing electrodes, while allowing the exchange of ions and metabolites needed for spontaneous electric activity (Fig. 1A,B). A dihydropyridine Ca^2+^ channel antagonist (nifedipine) was added to suppress smooth muscle contraction (Supplemental Fig. S2). The MEA detected field potential changes generated by the pacemaker activity of network-forming ICCs (Fig. 1C,D). This was confirmed by the MEA measurement in *W/W*^*v*^ mice that lack network-forming ICCs in the ileum (Supplemental Fig. S3).

The pacemaker potentials recorded by the 8 × 8 MEA were reconstructed into pseudo-color potential images (Fig. 1E) in which positive and negative field potentials, reflecting outward and inward currents, were assigned dark and light colors, respectively. Visualization of the MEA data enabled us to distinguish spatio-temporal features (Fig. 1F; Supplemental Fig. S1; also see Supplemental Videos SV1–4).

In the developmental process, pacemaker activities were divided into two patterns. ‘Expanding’ activity was defined by the initiation of activity in the MEA sensing area and a subsequent progressive expansion of the active region while the activity of the initiating region gradually subsided. As a result, ‘expanding’ activity typically formed an annular shape during the late phase (Expanding in Fig. 1F; Supplemental Video SV1). On the other hand, ‘migrating’ activity was defined by pacemaker activity migrating from a region adjacent to the MEA sensing area, and was manifested by an elongated active (light-colored) region that propagated approximately orthogonally to its long axis over the MEA sensing area in typical activities (Migrating in Fig. 1F; Supplemental Video SV2). Another characteristic feature of typical ‘migrating’ activity was a dark-colored area (indicating a transient positive potential) preceding the light-colored active area (further explanation is provided below).

A subclass of the micro-coordination pattern was defined by additional features during the propagating period of pacemaker events, independent of whether activity initiated in or migrated to the MEA sensing area. A ‘bumpy/aberrant’ event was due to aberrant spatio-temporal propagations resulting in a series of potential images lacking regularity, suggesting spatially confined electric coupling. In the example (Bumpy/Aberrant event in Fig. 1F; Supplemental Video SV3), spontaneous activity was initiated in the upper-left region and propagated toward the lower-right region while avoiding the lower-left region. Another subclass of pacemaker events was termed the ‘colliding/converging’ pattern, in which multiple propagating pacemaker potentials interacted in the MEA sensing area. Namely, they collided and/or converged to either cancel each other (Colliding in Fig. 1F; Supplemental Movie SV4) or merge and propagate in a different direction (explained later). This pattern of pacemaker event contained multiple activities which either initiated in or propagated to the MEA sensing area.

In 10 ileal muscle sheets obtained from wild-type mice, the frequency of pacemaker events ranged from 10–31 cycles per min (min^-1^), agreeing well with the values reported for ICCs in the myenteric plexus region (Kito and Suzuki, 2003). To verify the stability of the electric recording and biological conditions, features were compared between the first and second halves of the MEA measurement period (Fig. 1G). The frequency of pacemaker events and the occurrence of two basic activities displayed little change between the two halves (pacemaker event: 23.7 ± 5.6 vs. 25.2 ± 3.3 min^-1^ (23.6 vs. 13.1%), *P* = 0.234; expanding activity: 10.5 ± 9.7 vs. 12.7 ± 10.5 min^-1^ (92.3 vs. 82.7%), *P* = 0.307; migrating activity: 19.3 ± 14.1 vs. 18.2 ± 15.7 min^-1^ (73.1 vs. 86.3%), *P* = 0.376, *N* = 10). Of note, each pacemaker activity in ‘colliding/convergence’ and ‘bumpy/aberrant’ pacemaker events was summed as either one of the two basic activities (‘expanding’ or ‘migrating’ activity in Fig. 1H) by judging from the developmental process whether activity initiated in or migrated to the MEA sensing area.

### 3.2. Variations in the micro-coordination of pacemaker activity

Visualization of the MEA field potential data enabled us to realize considerably large variations of spatio-temporal pacemaker activity. Even in the same muscle sample, the initiating region of ‘expanding’ activity fluctuated in the MEA sensing area (Fig. 2A). In this sample, continuous recording demonstrated that most cases of propagating activity were of the ‘expanding’ type (12 out of 13 cycles), including the first three activities (in field potential traces in Fig. 2A). The first event was initiated in the upper-left region and propagated to adjacent regions (Expanding-1), the second event was initiated in the lower-right region (Expanding-2), and the third event was initiated near the center (Expanding-3) before expanding in an annular pattern (Supplemental Video SV5).

**Fig. 2.**
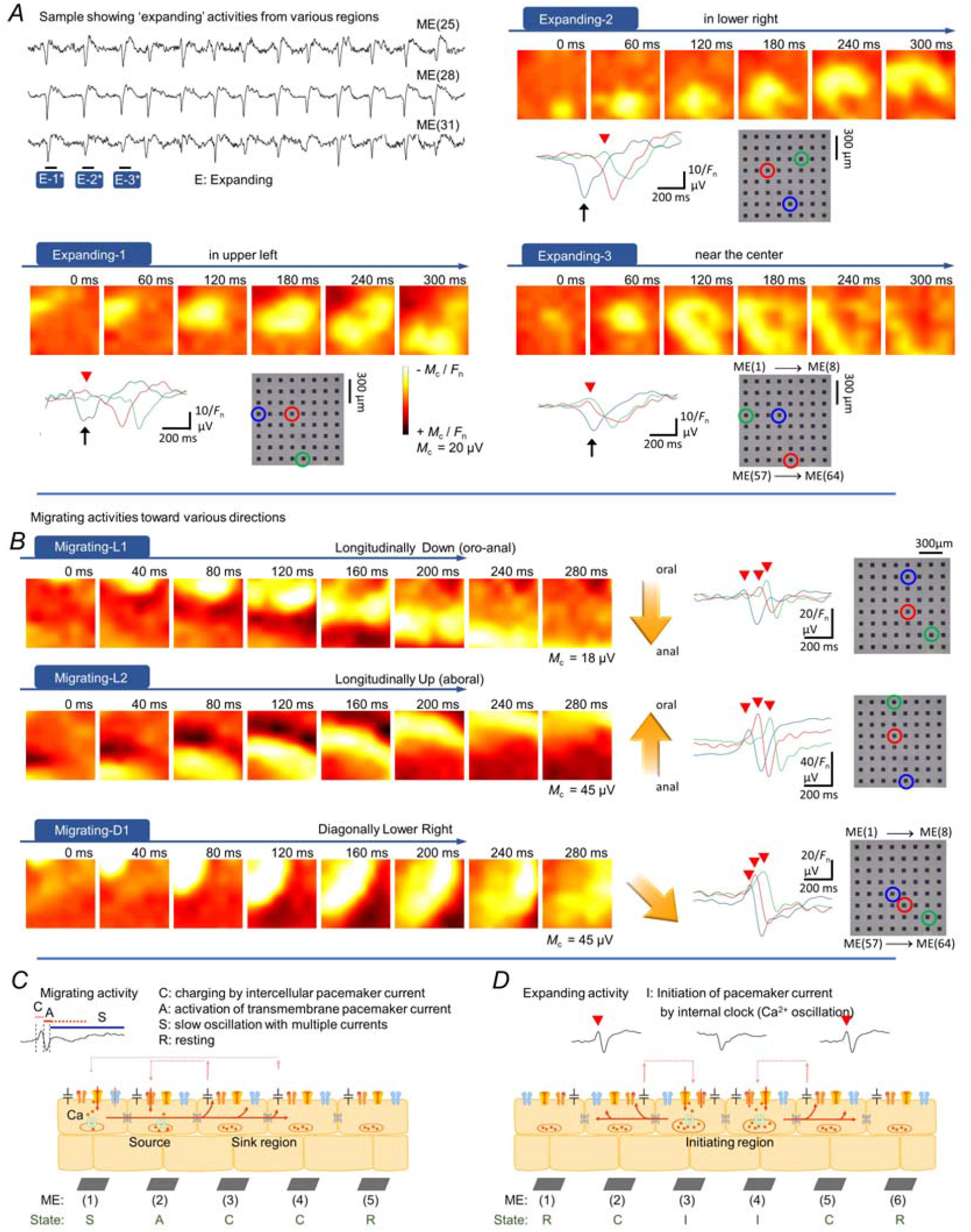
Variations of ‘expanding’ and ‘migrating’ activities (A,B), and schema showing cellular processes (C,D). (**A**) Field potentials recorded continuously by three microelectrodes [ME(25), ME(28) and ME(31)] arranged transversely on the same muscle preparation. The field potential images show three consecutive ‘expanding’ activities that were initiated in different regions: upper-left (Expanding-1), lower-right (Expanding-2) and near the center (Expanding-3), corresponding to E-1*, E-2* and E-3* in continuous field potential recording, respectively. M_c_: maximum potential used for color assignment. Traces of field potential waves recorded by three individual MEs are shown below each series of field potential images. Recorded MEs are indicated by the same color on the MEA. The electric potential wave near the initiating region (blue arrow) lacked a transient positive potential (red arrowhead). Representatives of ‘expanding’ activities observed in 10 samples. (**B**) Field potential images acquired from three samples, showing examples of ‘migrating’ activities propagating in different directions: longitudinally in the oro-anal direction (downward; Migrating-L1), longitudinally in the aboral direction (upward; Migrating-L2) and diagonally in the oro-anal direction (circumferentially leftward; Migrating-D1). Traces of representative field potential waves are shown along with each series of field potential images. Transient positive potentials (red arrowheads). Representatives of ‘migrating’ activities in 10 samples. (**C,D**) Illustrations showing the cellular processes proposed to underlie ‘migrating’ activity and ‘expanding’ activity, respectively (see text for an explanation). The square cells represent pacemaker interstitial cells along with adjacent smooth muscle cells electrically connected, thereby increasing electric capacitance.

‘Migrating’ activity exhibited variations in the direction of propagation (Fig. 2B). Notably, longitudinal propagation of activity in these two samples occurred in opposite directions, namely in the aboral (oral-to-anal, ordinally transporting) direction (Migrating-L1) and oral direction (Migrating-L2). In addition, pacemaker activity reversed in the circular (transverse) direction (Migrating-C1 and C2 in Supplemental Fig. S4), and propagated diagonally (Migrating-D1; Migrating-D2 in Supplemental Fig. S4). The propagation of ‘migrating’ activity in various directions rather than following the course of muscle layers suggests a major contribution of the ICC network (Fig. 1D) to the generation of this activity. We also noted that numerous features of propagation (i.e., amplitude, rate and direction) varied within the same muscle sample, even between two successive cycles of activity (Supplemental Video SV6).

Careful comparisons of potential images and field potential traces in ‘expanding’ activity indicated that the pacemaker potential lacked a preceding transient positive potential in the initiating microregion (blue arrow in Fig. 2A). On the other hand, in ‘migrating’ activity, a dark-colored area preceded a light-colored area when activity propagated from the adjacent area of the MEA sensing area (Fig. 2B; Supplemental Fig. 1G, H). This corresponds to a transient positive potential (red arrowhead) preceding the fast down-stroke of the pacemaker potential.

The processes of propagating activity in the MEA sensing area were tentatively modeled as follows. In ‘migrating’ activity, the light- and dark-colored areas are considered to be the ‘source’ and ‘sink’ of a local circuit current, respectively (see schematic illustrations in Fig. 2C and Supplemental Fig. S3). The intercellular pacemaker current progressively charges the membrane in the sink region until the transmembrane pacemaker current is activated, employing a voltage-gated mechanism (Goto et al. 2004). Thereby, propagation of pacemaker activity sequentially along the muscle sheet acts as a ‘volume conductor’ (Supplemental Fig. S5). On the other hand, a cellular model of ‘expanding’ activity was constructed by adding an initiating region which likely employs an internal clock mechanism such as the activation of Ca^2+^-activated transmembrane pacemaker current in response to intracellular Ca^2+^ oscillations owing to Ca^2+^ release channels in the endoplasmic reticulum (ER) (Fig. 2D; Supplemental Fig. S6) (Aoyama et al. 2004; Saeki et al. 2019). After ‘expanding’ activity has been initiated, it can subsequently propagate away from the initiating region by the same mechanism as ‘migrating’ activity. However, some of the pacemaker potentials near the initiating site did not have preceding positive potentials (Fig. 2A), implying that propagation in the immediate vicinity of the initiating region may contain pacemaker cells coincidently activated by an internal Ca^2+^ clock, but with some delay. An alternative possibility is that a small group of cells neighboring the initiating region were activated by the intercellular propagation of Ca^2+^ or intracellular messengers (e.g., IP_3_) via gap junctions, which has been reported for other cell types (Halidi et al. 2011).

### 3.3. Interconversion of micro-coordination patterns

When pacemaker events were classified into four patterns by incorporating the features in a propagation period, in addition to the large variations within the same category of event, we also frequently observed interconversion between the patterns (eight out of ten samples during the continuous MEA measurement). The frequency of interconversion differed substantially between samples. Continuous MEA recordings from a muscle sheet with frequent interconversions exhibited eight consecutive pacemaker events: one ‘migrating’, four ‘expanding’ (with different initiating regions), two ‘colliding’, and one ‘bumpy/aberrant’ (Fig. 3A). In the ‘migrating’ event, activity crossed the MEA area in a diagonal direction (Fig. 3B), and in the ‘expanding’ event, activity was initiated in the upper-left region (Fig. 3C; the last ‘expanding’ event in Fig. 3A). In the ‘bumpy/aberrant’ event, activity initially propagated from the lower region of the image in an aboral direction, but the activity only reached the upper-left region and failed to propagate to the upper-right region (Fig. 3D). In the first of the two ‘colliding’ events (Colliding-1: Fig. 3E), two pacemaker activities expanded from the upper-left and lower-middle regions to collide near the center, following which the merged activity propagated toward the upper-right region. In the second ‘colliding’ event (Colliding-2: Fig. 3F), two pacemaker activities initially propagated similar to the first ‘colliding’ event, but the merged activity did not propagate toward the middle-right region showing a slowly oscillating potential (Fig. 3F). It is likely that the aberrant slow potential substantially prolonged the refractory period of pacemaker cells in this region, resulting in a ‘bumpy/aberrant’ event. Indeed, various types of regional aberrant electric activity occurred, which may account for ‘bumpy/aberrant’ events. For example, in another sample showing an interconversion of micro-coordination patterns, multiple pacemaker activities occurred independently in the MEA sensing area and were separated by a ‘non-conducting’ region where rapid potentials run frequently (Supplemental Fig. S7).

**Fig. 3.**
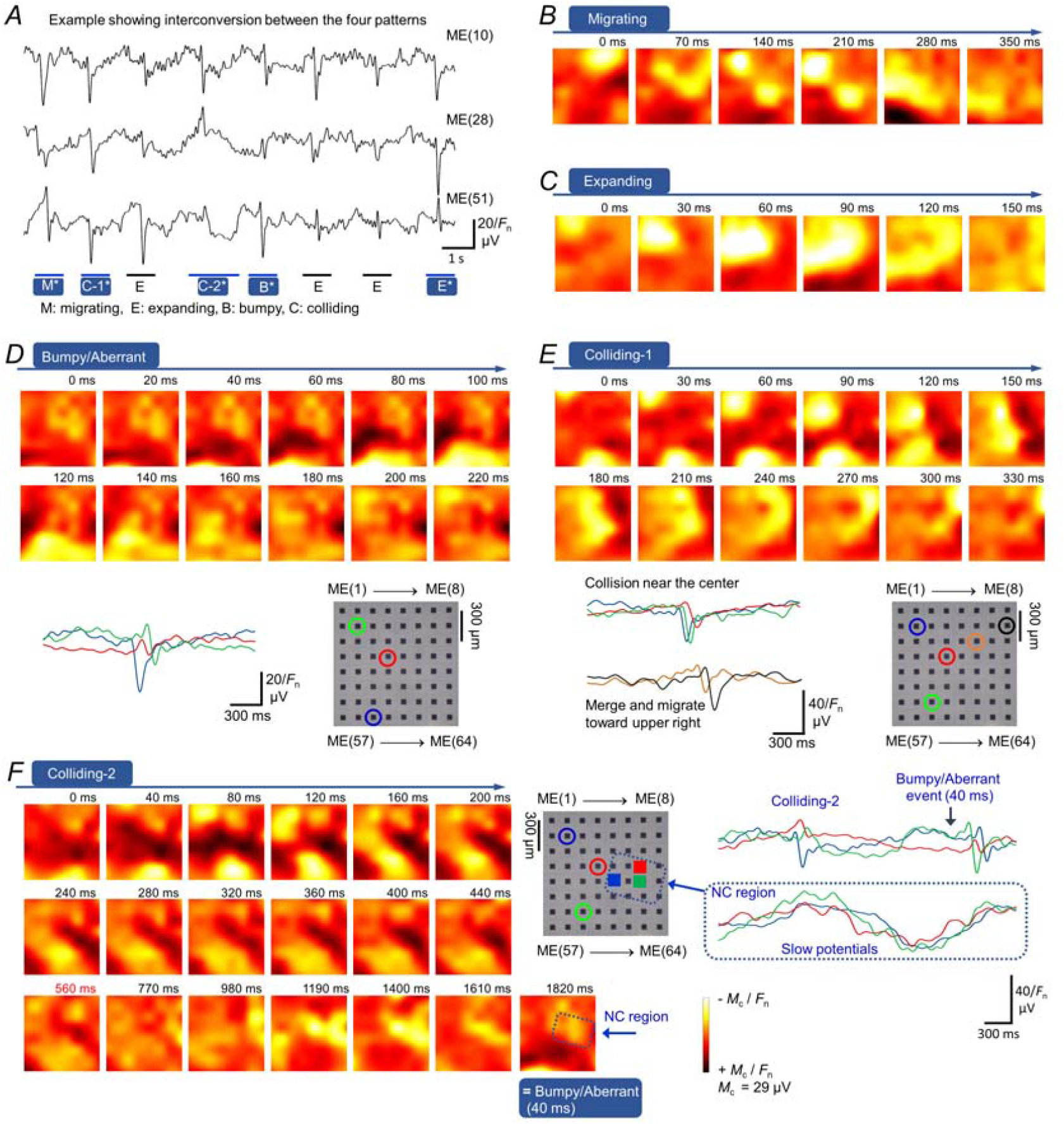
Interconversion of micro-coordination patterns. (**A**) Field potentials continuously recorded from a muscle sample by three microelectrodes [ME(10), ME(28) and ME(51)], showing interconversion between the four micro-coordination patterns of pacemaker events. This sample is a representative of the 10 samples used in Fig. 1 (G, H). Asterisks indicate the pacemaker events shown as potential maps in **B–F**. (**B**) Potential images of the ‘migrating’ event (activity) shown in **A** (M*). (**C**) Potential images of the ‘expanding’ event (activity) shown in **A** (E*). (**D**) Potential images of the ‘bumpy/aberrant’ event shown in **A** (B*) together with superimposed field potential traces recorded by different microelectrodes. (**E**) Potential images of the first ‘colliding’ event shown in **A** (C-1*) together with superimposed field potential traces recorded by different microelectrodes. (**F**) Potential images of the second ‘colliding’ activity shown in **A** (C-2*) and the phase that followed. The superimposed field potentials reveal slow potentials in the middle-right region (‘bumpy’ spot), suggesting that this may have caused a ‘bumpy/aberrant’ event in the next cycle of pacemaker activity.

### 3.4. Proof-of-concept through the action of 5-HT

The micro-coordination pattern of pacemaker activity was changeable (Fig. 3), but the frequency of occurrence of each pattern was relatively stable in the control condition (Fig. 1**H,I**). We thus attempted to examine if their frequencies could be employed for functional assessment in a digital manner, like the ‘open’ and ‘closed’ states of ionic channels in patch clamp experiments. Since 5-HT is one of the most important signaling molecules in functional modulation of GI motility (Spiller, 2011; Mawe and Hoffman, 2013), we examined the effects of 5-HT after each control MEA measurement (Fig. 4). An example of a series of MEA measurements from a muscle sheet demonstrated frequent activities of the ‘expanding’ pattern: a typical cycle involved the initiation of activity in the upper-middle region in control conditions (Fig. 4A,B; Supplemental Video SV7), and those converted into a ‘migrating’ pattern in the presence of 5-HT (100 μM), accompanied by an increase in the frequency of pacemaker events (Fig. 4C,D; Supplemental Video SV8). Similar changes were observed in nine muscle sheets (see also Supplemental Fig. S8: another example of the consistent effects of 5-HT examined in a muscle sheet showing frequent ‘bumpy/aberrant’ events in the control).

**Fig. 4.**
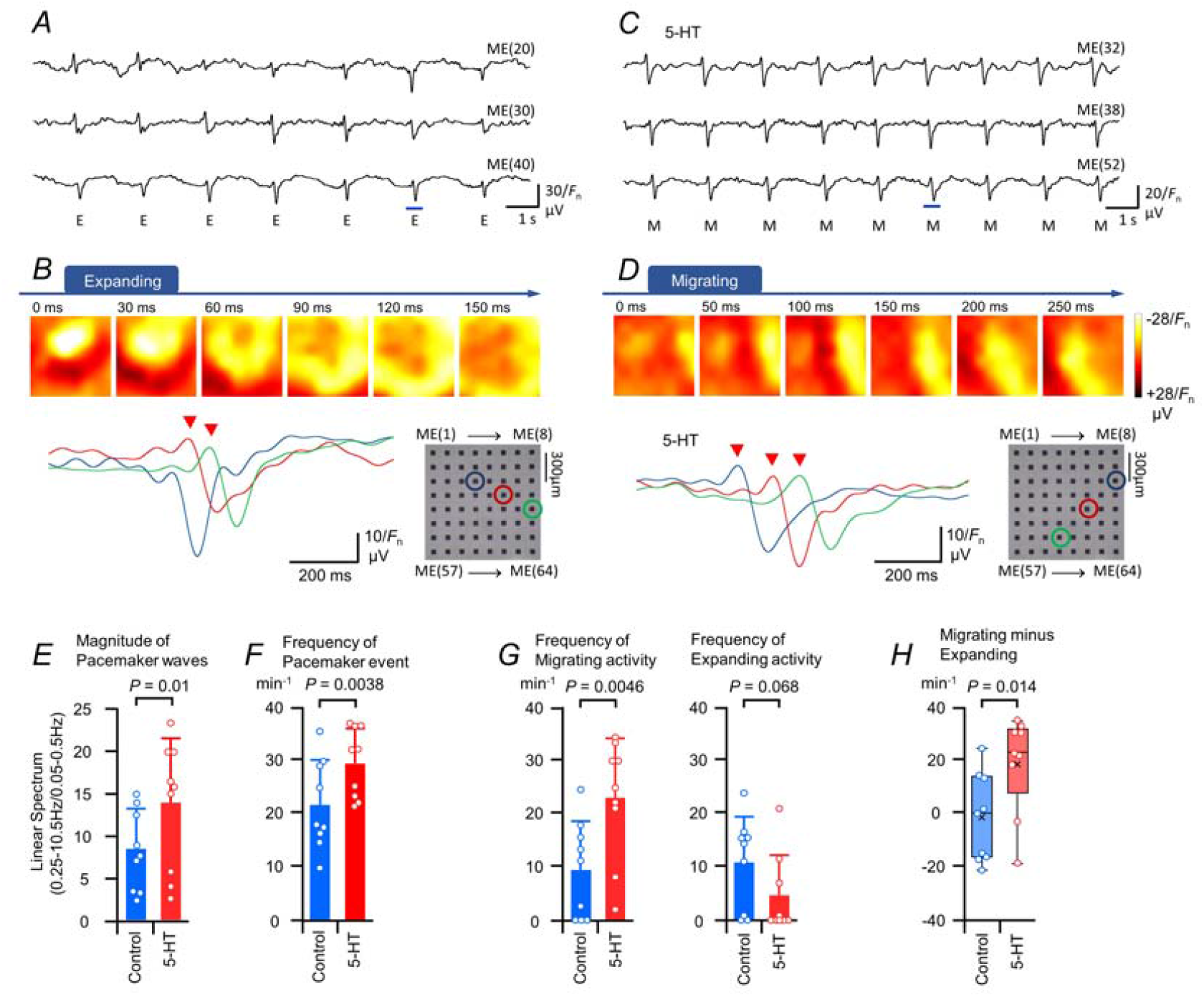
Modulation of micro-coordination patterns by 5-HT. (**A**) Field potentials continuously recorded in a sample of ileal musculature in control medium. All activities are of the ‘expanding’ type. (**B**) Series of field potential images and superimposed field potential traces for the period marked with a bar in A. (**C, D**) Effects of 5-HT (100 μM) on pacemaker activity in the same sample. All activities are of the ‘migrating’ type. The size of the potential image is 1050 × 1050 μm^2^. (**E, F**) Graphs showing the ratio of the normalized magnitude of pacemaker waves (E: LS_0.25–10.5Hz_/LS_0.05-0.5Hz_) and the frequency (F: min^-1^) of pacemaker events under control conditions and in the presence of 5-HT. (**G**) Graphs showing the frequency of ‘migrating’ or ‘expanding’ activity (min^-1^). (**H**) Box-and-whisker plot of the difference between the frequency of migrating and expanding activity (min^-1^) in **H**. ×, the mean; vertical line, the median value. (**E-H**, control vs 5-HT in the same sample: nine samples from nine mice).

Linear spectrum (LS) analysis revealed that the magnitude of the pacemaker potential imaging frequency range (LS_0.25–10.5Hz_) normalized whereas that of the sub-basal frequency range (LS_0.05-0.5Hz_) increased in the presence of 5-HT (LS_0.25-10.5_/LS_0.05-0.5_: 8.3 ± 4.7 in the control vs. 13.7 ± 7.5 in 5-HT (56.6 vs. 54.7%), *P* = 0.01, *N* = 9; Fig. 4E). On the other hand, the frequency of pacemaker events increased significantly in the presence of 5-HT (27.3 ± 7.2 min^-1^ in 5-HT vs. 21.3 ± 8.4 min^-1^ in the control (26.4 vs. 39.4%), *P* = 0.0038, *N* = 9; Fig. 4F). In addition, 5-HT increased the occurrence of ‘migrating’ activity from 9.4 ± 9.1 to 23.0 ± 11.1 min^-1^ (96.8 vs. 48.2%, *P* = 0.0047, *N* = 9; Fig. 4G), and decreased that of ‘expanding’ activity from 10.9 ± 8.6 to 4.6 ± 7.4 min^-1^ (78.9 vs. 161%, *P* = 0.068, *N* = 9; Fig. 4G) with the difference between the frequency of migrating and expanding activity being –1.48 ± 16.4 in the control vs. 18.4 ± 18.0 in 5-HT (1108 vs. 97.8%, *P* = 0.014, *N*=9, Fig. 4H). In the sum of nine experiments (total *N* = 9, 165 events in the control vs 216 events in the 5-HT treatment for 54 s), the number of the four pacemaker events changed accordingly. ‘Expanding’ and ‘bumpy/aberrant’ events decreased (from 49 to 10 events and from 35 to 20 events, respectively) while ‘migrating’ and ‘colliding/converging’ events increased (from 69 to 134 events and from 12 to 52 events, respectively).

### 3.5. Biological significance of the two major patterns

As the width of sensing electrode becomes small, the transmission efficiency of low frequency electric signals decreases due to an increase in impedance. Pt black nanoparticles have resolved electrical contradictions by increasing the surface area of sensing microelectrodes (Taniguchi et al. 2013; Iwata et al. 2017). Thereby, the dialysis membrane-reinforced low-impedance MEA technique enabled the stable measurement of field potentials in a wide frequency range, and the classification of spatio-temporal coordination of pacemaker potentials in a sensing microregion, despite surprisingly large variations. We note that the electrical matching of the sensing microelectrode and detecting amplifier system is important to appropriately measure low-frequency field potentials, which is a characteristic feature of the GI tract.

To date, evidence has accumulated for the synchronization of electric potentials and intracellular Ca^2+^ oscillations in gut pacemaker activity (Torihashi et al. 2002; van Helden and Imtiaz, 2003; Park et al. 2006). Therefore, the initiation of a pacemaker potential, such as ‘expanding’ activity in the MEA sensing microregion, is considered to reflect the Ca^2+^ clock mechanism (Fig. 2D and Supplemental Fig. S6) operated by the co-existence of ryanodine receptors and inositol 1,4,5-trisphosphate receptors (IP_3_Rs) in the ER of ICCs (Aoyama et al. 2004; Saeki et al. 2019) that consequently activate transmembrane Ca^2+^-activated Cl^-^ conductance (Dickens et al. 1999; Hwang et al. 2009). On the other hand, ‘migrating’ activity, defined by activity that propagates to the MEA sensing area, represents pacemaker currents that employ voltage-gated mechanisms.

Typical ‘migrating’ activity was characterized by a dark-colored (positive potential) area at the front of the propagating waveform that was followed by a light-colored (negative potential) area. The spatiotemporal relationship between the light- and dark-colored areas implies that intercellular coupling causes the ICCs in the dark-colored region to become electrically charged by the intercellular pacemaker current that has been activated in the ICCs in the light-colored region (sink and source regions, respectively, in Fig. 2C). In other words, pacemaker activity propagates through a volume conductor due to the formation of local circuits. This feature explains the existence of voltage-gated inward current attributed to atypical non-selective cation channels (Goto et al. 2004) and TTX-resistant Na^+^ channels (Na_v_1.5) (Strege et al. 2005) in pacemaker cells of rodents and humans, respectively. Taken together, the image analysis in the present study thus validates the biological significance of duplicating pacemaker mechanisms in the small intestine, by implying the functional state of the sensing microregion (i.e., spontaneity and synchrony/continuity for the ‘expanding’ and ‘migrating’ patterns, respectively).

### 3.6. Complexity of the waveform in extracellular recording

The potential images explicitly demonstrate the two sources of pacemaker current via transmembrane and intercellular pathways that can account for the complex waveform of electric activity in extracellular recordings (Sanders et al. 2016). Namely, during the depolarization period, the directions of transmembrane and intercellular currents were opposite: Transmembrane current flows inwardly while intercellular current flows outwardly to charge the membrane as a volume conductor propagates [e.g. microregions near ME(3) and ME(2), respectively, in Fig. 2C]. Extracellular electrodes detect changes in field potential, thereby reflecting the direction of these currents, when the spatial resolution is sufficient to detect the propagation of volume conductor. On the other hand, intracellular electrodes only exhibit depolarization when detecting either transmembrane or intercellular currents, so the first-derivative of the (trans)membrane potential waveform differs from the extracellular field potential waveform in the electric syncytium. The waveform is further complicated in microregions where electric potentials from multiple sources interact (e.g., Fig. 3F, Supplemental Fig. S7). In ‘migrating’ pacemaker activities recorded in the present study, therefore, the fact that preceding positive potentials (outward current) of sizeable amplitude propagate in the MEA sensing area, verifies the electric syncytium nature of the GI tract (Tomita, 1970; Manchanda et al. 2019), where smooth muscle cells and network-forming pacemaker cells are electrically connected (Dickens et al. 1999; Sperelakis and Daniel, 2004).

The present field potential microimaging revealed that the direction of ‘migrating’ activity varied considerably. Actually, volume conductors flexibly migrate, irrespective of the running circular and longitudinal muscle bundles (e.g., Fig. 2B; Supplemental Fig. S4), indicating that pacemaker currents conduct mainly via a network of ICCs (i.e., ICCs in the myenteric plexus region). The lack of pacemaker potentials in the ileum of *W/W*^*v*^ mice reinforces this hypothesis. The use of an extracellular electrode array with a wide polar distance (1 mm) and recording area (10 × 10 mm^2^) was previously reported for the existence of reentrant electric waves in the small intestine of healthy rats, but they were less frequent than in diabetic rats (Lammers et al. 2011, 2012). The circuit that leads reentrant electric waves may correspond to large variations of ‘migrating’ activity observed in the small MEA area of this study. However, measurements were performed in different conditions of extracellular medium and preparation. For instance, previous measurements of electrode array with a wide polar distance were conducted without using Ca^2+^ channel antagonists which suppress smooth muscle electric activity (Lammers et al. 2011, 2012), while our MEA measurements used musculature preparations in which the influence of sensory nerves and humoral factors, such as 5-HT and incretins were isolated from the mucous. Therefore, careful assessment is needed to compare the spatio-temporal features of spontaneous electric activity between studies. In addition, these previous studies probably did not distinguish whether activity ‘migrates’ to or ‘expands’ in the recording region, because the size of the volume conductor (i.e., propagating as coupled dark and light regions in Fig. 2B) is comparable to or smaller than the area covered by each sensing electrode (1 × 1 mm^2^). In other words, the coupling mechanism between electric waves is obscure due to either electric conduction or an internal clock of the relaxation oscillator. It is likely that the total electric dynamics in a large area of the small intestine consists of a mixture (i.e., patchwork) of microregions with ‘migrating’ (electrically conducting) and ‘expanding’ (initiating) activities.

### 3.7. Alteration of micro-coordination patterns related to motor function

5-HT signaling is involved in promoting the propulsion of foodstuff through the intestine (Bülbring and Lin, 1958; Mawe and Hoffman, 2013). Therefore, during 5-HT application, an increase in the ratio of ‘migrating’ to ‘expanding’ activity (Fig. 4) likely reflects a change in the microregion from a segmentation-preferable state to a propulsion-preferable state. In addition, it is considered that the decrease in ‘bumpy/aberrant’ events and increase in ‘colliding/converging’ events indicate improved electric connectivity between network-forming myenteric ICCs. In turn, this allows pacemaker activity originating in areas adjacent to the MEA to reach the MEA-sensing area more frequently. Consistently, the ratio of ‘migrating’ to ‘expanding’ activity (R_M/E_) implies a three-fold enlargement of the area covered by each pacemaker activity initiating site (i.e., R_M/E_ + 1: ~1.9 in the control vs. ~6.4 in 5-HT, see Methods).

In the small intestine, it was reported that aberrant slow electric oscillations originating from different cellular sources, such as ICCs-DMP, yield segmentation of the tract by waxing and waning the basal pacemaker activity in network-forming myenteric ICCs (Huizinga et al. 2014). It is speculated that 5-HT might act to amplify pacemaker activity by suppressing such aberrant spontaneous rhythmicity of low frequency originating from cell members other than network-forming ICCs-MP (e.g., slow potentials as shown in Fig. 3F) thereby improving local electric conduction in microregions. This hypothesis agrees well with the fact that the magnitude of pacemaker waves relative to that of sub-basal waves increased in the presence of 5-HT, mainly reflecting the latter component of field potential waves (Supplemental Fig. S9). In human genetics, mutations of 5-HT-related genes are involved in patients with GI dysmotility, such as the LL variant of the serotonin-transporter-gene-linked polymorphic region (5-HTTLPR) for constipation-predominant IBS (Zhang et al. 2014), or the HTR3E-untranslated region variant for female diarrhea-predominant IBS (Kapeller et al. 2008). We anticipate that the MEA method communicated in this study will be used for investigating spatio-temporal properties of pacemaker potentials in small model animals with bowel motility diseases, especially those associated with 5-HT signaling and pacemaker activity.

## 4. Conclusion

The dialysis membrane-reinforced low-impedance MEA technique enabled the stable measurement of field potentials in a wide frequency range in the microregion of isolated musculature by resolving electrical contradictions. In this study, we analysed pacemaker dynamics in the small intestine of the mouse to assess the possible role of the network-forming pacemaker cells in coordinated GI movement. Visualization of the micro-coordination of pacemaker potentials allowed them to be classified into several basic patterns, despite huge variations. The two major ‘migrating’ and ‘expanding’ patterns were considered to reflect duplicating pacemaker current systems that employ voltage-gated and Ca^2+^ oscillation-activated channels, respectively (Fig. 2C,D). Of note, during the propagation of pacemaker potentials, our field potential microimaging exhibited volume conductors as coupled dark and light regions that correspond to positive intercellular and negative transmembrane currents, respectively, although both charged the plasma membrane positively. The opposite directions of the intercellular and transmembrane currents accounted for the complex waveforms of extracellular recording.

The pattern of pacemaker micro-coordination was changeable in each event, but the ratio of patterns was similar in the same microregion of samples during a sufficiently long period (Fig. 1G-I). This study revealed that 5-HT significantly increased the occurrence of ‘migrating’ pattern of activity (Fig. 4), agreeing well with its action, which causes a shift in the contractile function of the GI tract from segmentation to propulsive motions. These results therefore provide evidence that despite the requirement of further refinement, our MEA method of microimaging is beneficial for quantitative assessment of electric coordination underlying GI motility in health and disease of small animals.

## CReDiT Author contributions

NI: Investigation, Data curation, Visualization, Validation, Writing – review & editing.

CT: Investigation, Data curation, Visualization.

MY: Investigation, Data curation.

NM: Investigation, Data curation, Writing – review & editing.

NK: Investigation, Data curation.

YK: Conceptualization, Validation, Writing – review & editing.

SN: Conceptualization, Methodology, Investigation, Data curation, Validation, Funding acquisition, Writing – original draft.

## Acknowledgments

We are grateful to Tatsuhiro Noda, Hirotaka Morishita and Tomoka Nomura (Department of Cell Physiology, Nagoya University Graduate School of Medicine) for their assistance with the MEA experiments.

This work was partly supported by a grant-in-aid for Scientific Research from the Japan Society for the Promotion of Science (JSPS No. 19H03558) and a research grant from the Suzuken Memorial Foundation.

## Data Availability

All data supporting the results presented in the manuscript are included in the manuscript figures (n≤10).

## Legends for Supplemental Figures

**Fig. S1.**
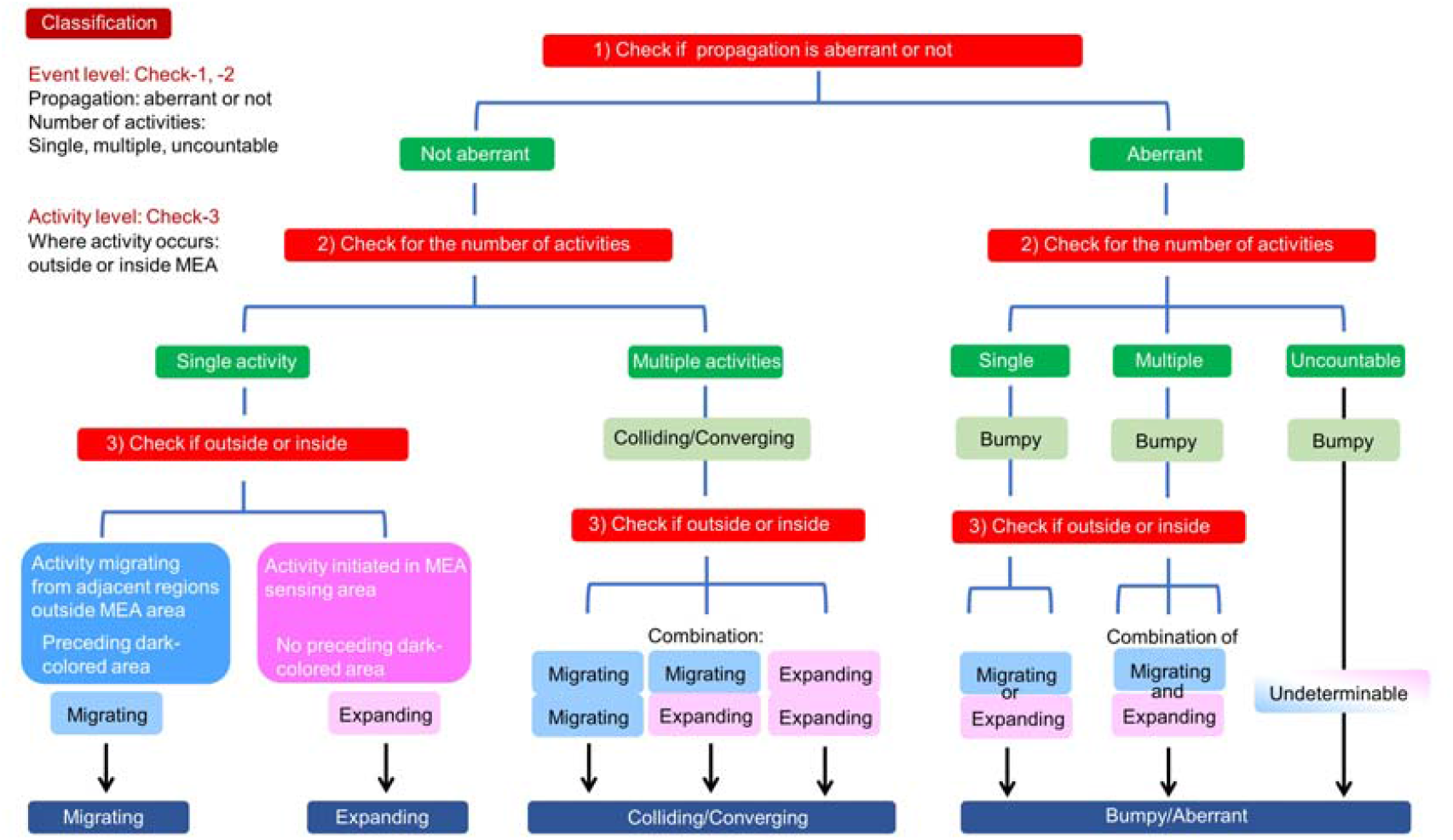
Illustration showing the procedures of classification in micro-coordination of pacemaker potentials. For this classification, there are three check points in event and activity levels. (Check-1) At first, each pacemaker event (pacemaker period) was classified by the propagation of pacemaker potentials in a coherent or incoherent/aberrant manner. The latter pacemaker event was classified into a ‘bumpy/aberrant’ event. (Check-2) Secondly, the pacemaker event was classified by the number of pacemaker activities that independently occurred. (Check-3) Thirdly, each pacemaker activity was classified into a ‘migrating’ or ‘expanding’ activity pattern by the initiation process (i.e., where activity occurs, outside or inside, respectively, the MEA sensing area). In a pacemaker event containing single pacemaker activity, ‘migrating’ and ‘expanding’ activities correspond to ‘migrating’ and ‘expanding’ events, respectively. In a ‘bumpy/aberrant’ event, if the number of activities is not countable, the pattern of activity (‘migrating’ or ‘expanding’) cannot be determined. The case of an uncountable, ‘bumpy/aberrant’ event was not used in this study.

**Fig. S2.**
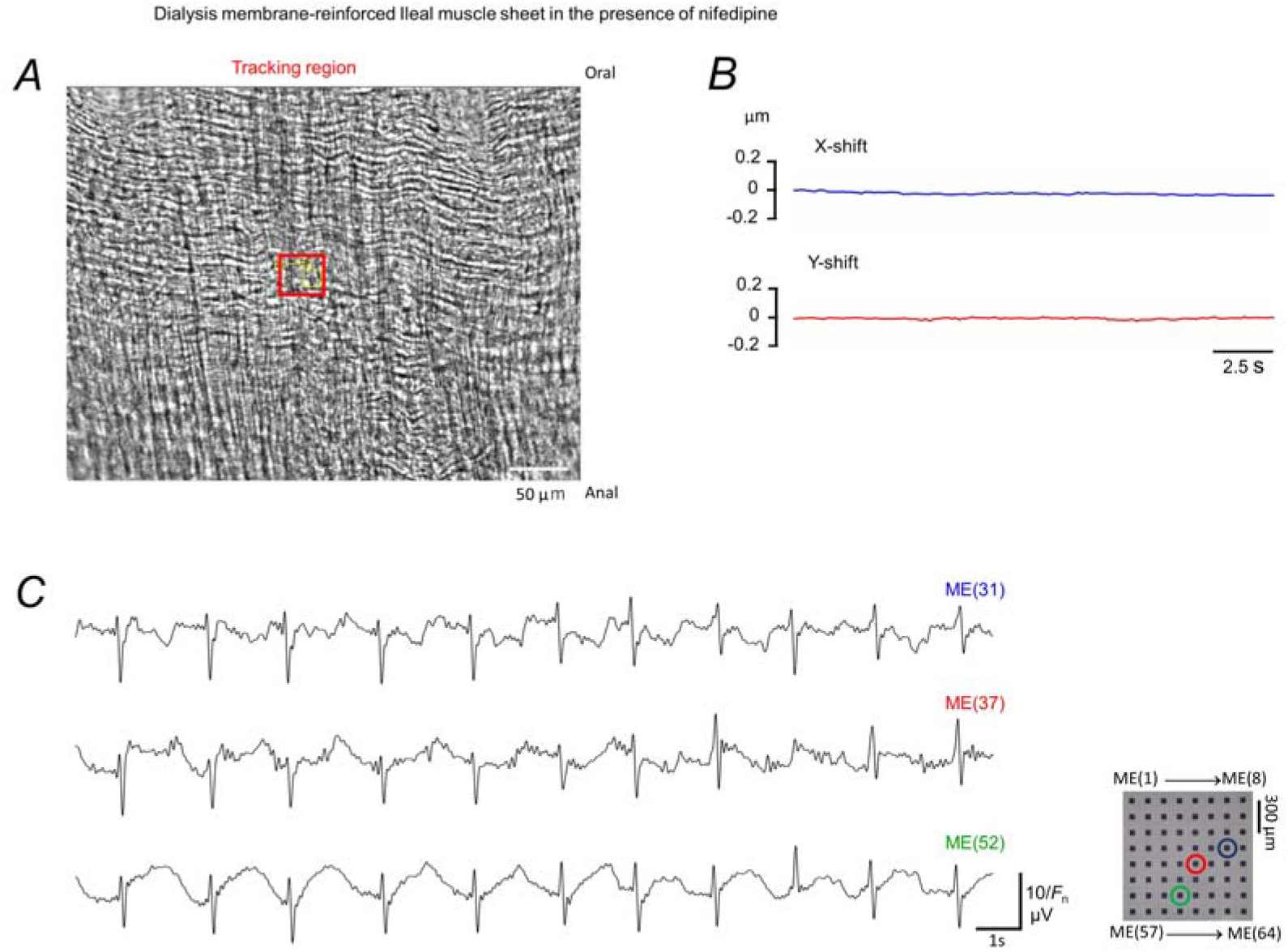
Motion tracking of a muscle sample used in MEA measurement. (**A**) After the MEA measurement, video of transmission image acquired in a muscle sample mounted in a glass-bottomed chamber. The extracellular solution contained nifedipine. The chamber was warmed on a glass heater. (**B**) Plots of the X, Y-shift of the tracking square shown in **A**. (**C**) Plots of representative field potentials of the MEA. The measurement was performed before the video, verifying negligible movement.

**Fig. S3.**
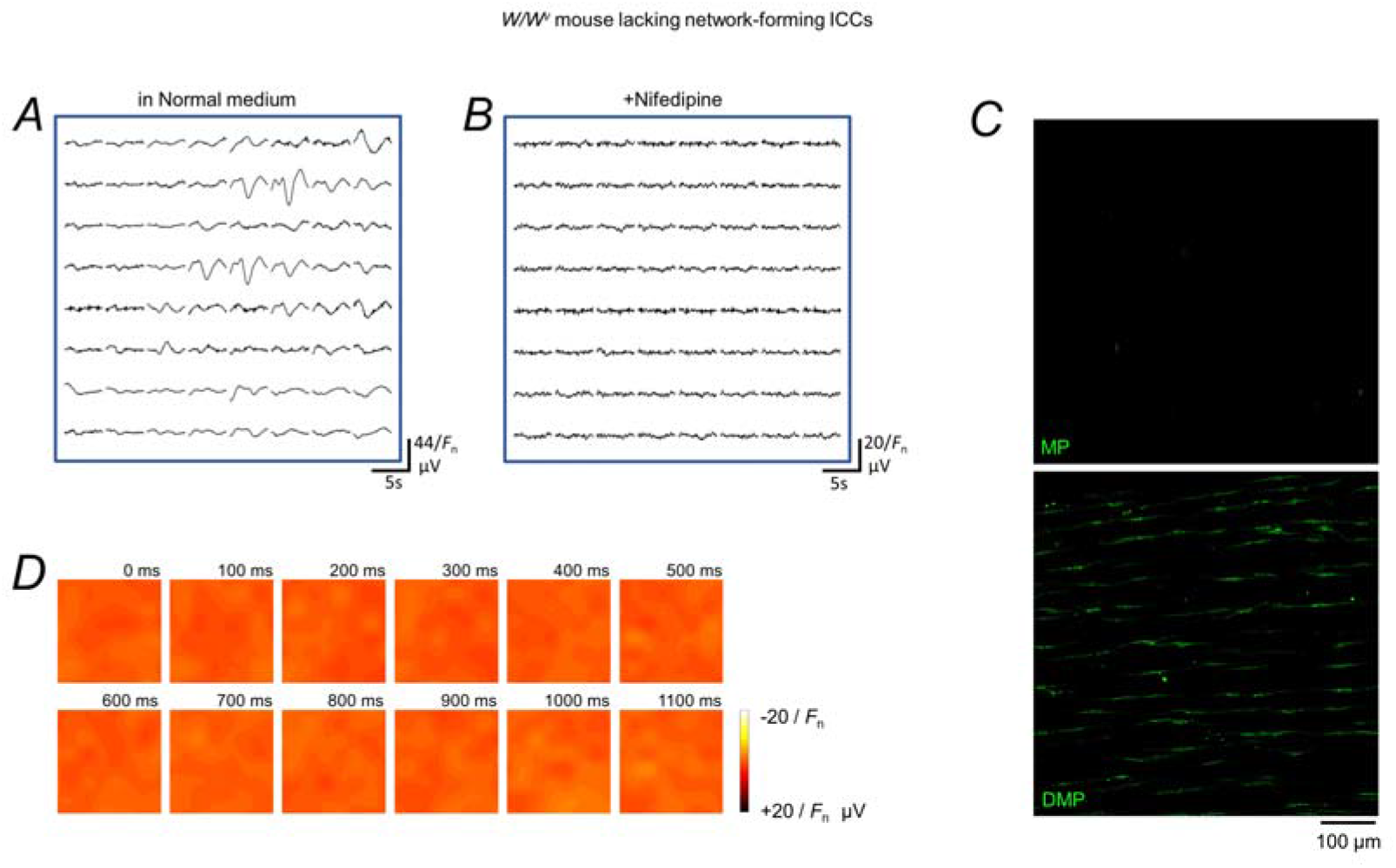
Microelectrode array (MEA) recordings of spontaneous electrical activity in a sample of ileal muscle from a *W/W*^*v*^ mouse that lacks network-forming interstitial cells of Cajal (ICCs). After removal of the mucosa, the ileal muscle sample was mounted on an 8 × 8 MEA using a piece of dialysis membrane, as shown in Fig. 1A. (**A, B**) An 8 × 8 plot of field potentials recorded under control conditions (**A**) and after exposure to the dihydropyridine Ca^2+^ channel antagonist, nifedipine (**B**). (**C**) Confocal images of c-kit immunoreactivity in the regions of the myenteric plexus (upper) and the deep muscular plexus (lower). (**D**) Field potential images showing electrical activity in the presence of nifedipine, corresponding to the MEA measurement in (**B**). In contrast to the MEA recordings made in wild-type mice, there was negligible electric activity in the ileum of *W/W*^*v*^ mice in the presence of nifedipine (**B**).

**Fig. S4.**
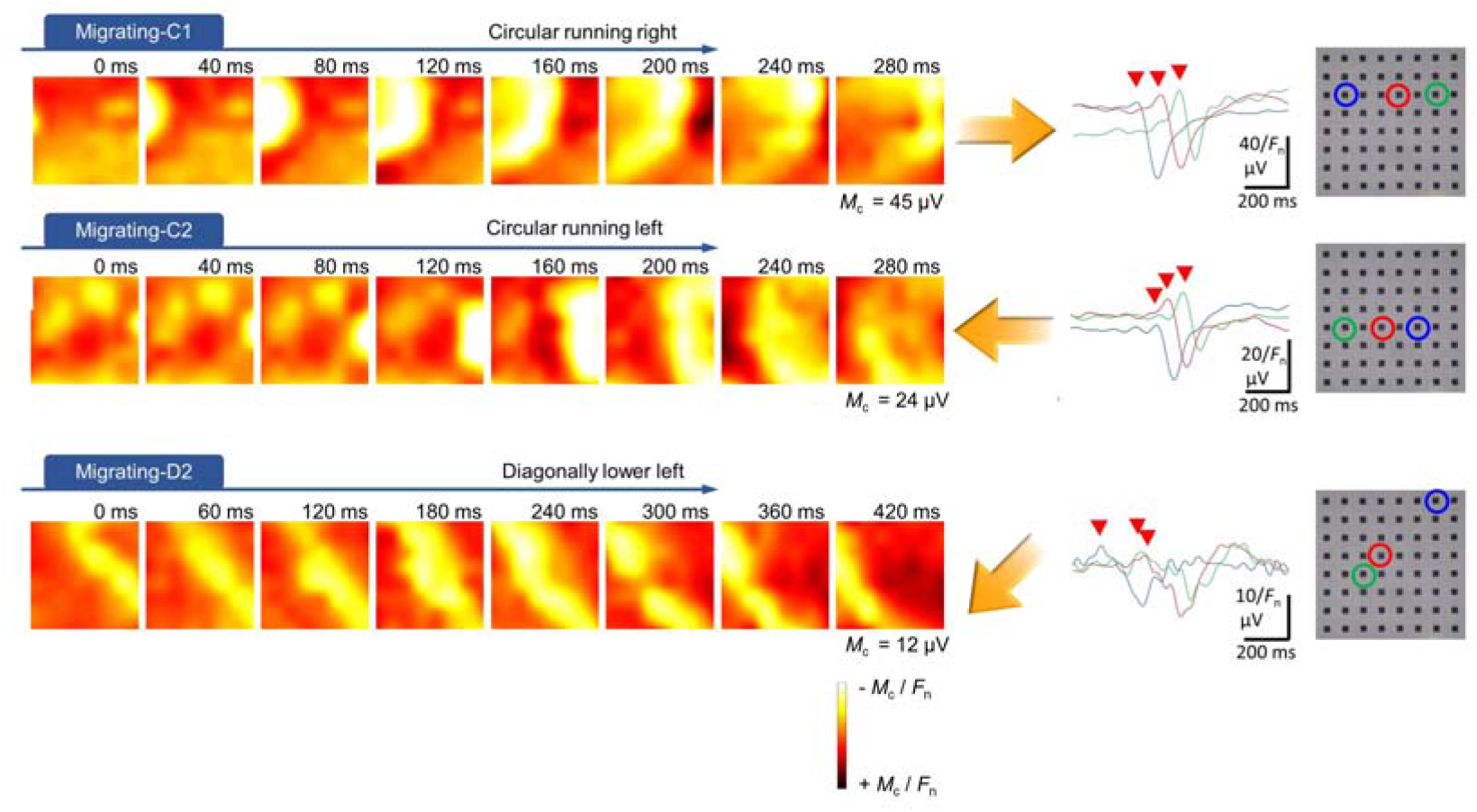
‘Migrating’ activities propagated in circular and diagonal directions. Field potential images and representative field potential waves showing three examples of ‘migrating’ activity, in addition to those in Fig. 2B. Migrating-C1 and -C2: Circumferential propagation (along the circular muscle) toward the right and left, respectively. Migrating-D2: Diagonal propagation in the oro-anal direction (circumferentially toward the left). M_c_: maximum potential used for color assignment. Red arrowheads indicate transient positive potentials in the ‘migrating’ waves. The MEs for representative field potential waves are indicated by circles of the same color on the MEA.

**Fig. S5.**
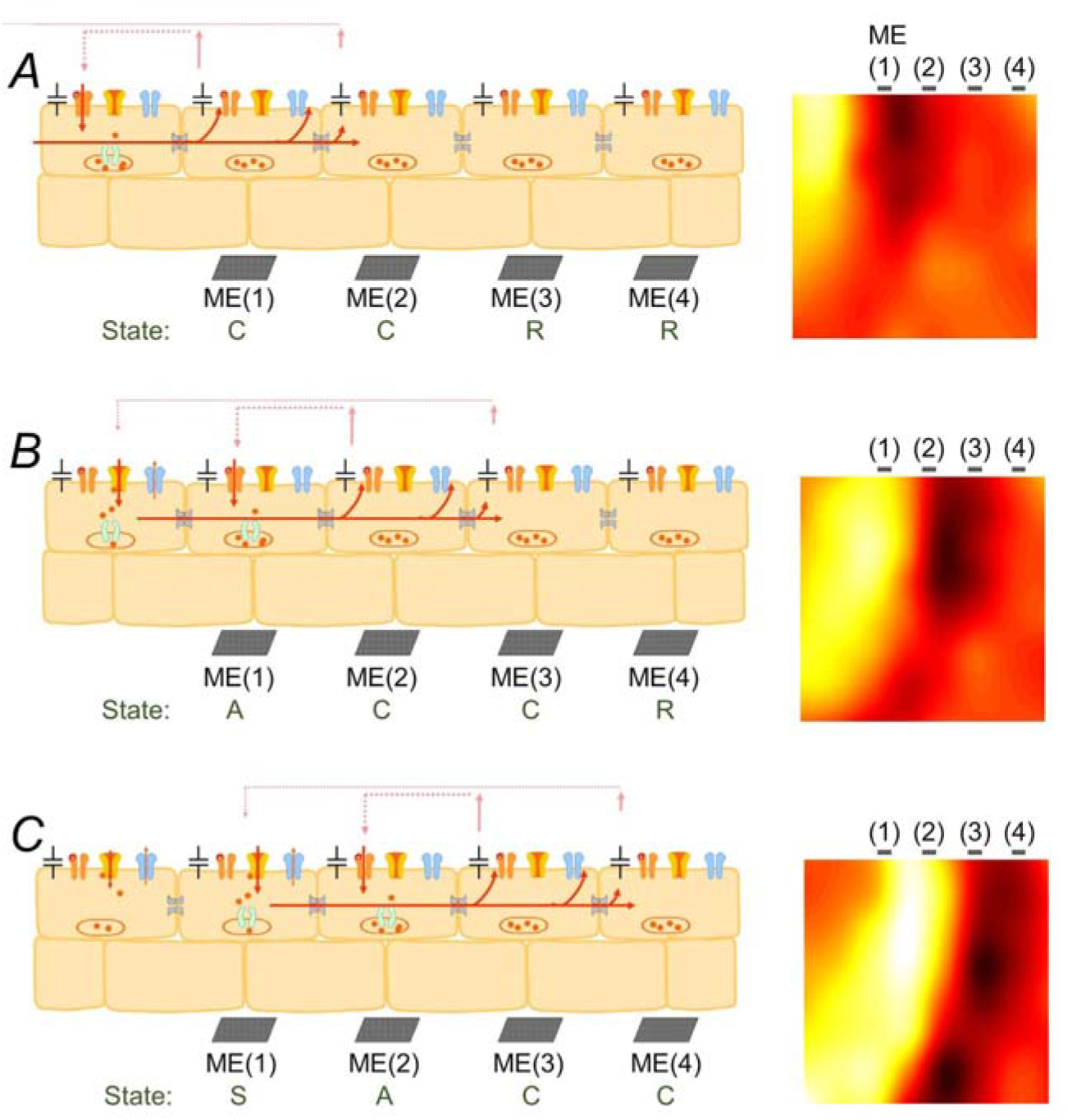
Schematic illustration showing the process underlying ‘migrating’ activity. (**A**) Intercellular pacemaker current propagates toward the cells near ME(1) and ME(2) and charges their plasma membranes. The cells near ME(1) and ME(2) are in a charging (‘C’) state, and those near the other MEs are in a resting (‘R’) state. (**B**) Next, intercellular pacemaker current depolarizes the plasma membrane to the threshold and activates voltage-gated inward current (a part of pacemaker current) in the cells near ME(1) so that these cells enter an activated (‘A’) state. Meanwhile, the intercellular pacemaker current propagates toward the cells near ME(2) and ME(3). (**C**) The cells near ME(1) enter a slow oscillation (‘S’) state in which transmembrane outward K^+^ current and Ca^2+^-activated inward current are continuously activated. Intracellular Ca^2+^ release processes likely contribute to the activation of Ca^2+^-activated inward currents (e.g., anoctamin-1 Ca^2+^-activated Cl^-^ channels: the equilibrium potential of pacemaker cells ≈ -20 mV, more positive than the resting membrane potential). Meanwhile, the cells near ME(2) enter the ‘A’ state, and the intercellular pacemaker current further propagates toward the cells near ME(3) and ME(3). Thus, the cells above ME(1) to ME(4) form a local circuit. The intercellular pacemaker current does not propagate toward cells on the left, because the membranes of these cells are already charged. Each cell in the scheme represents a group of electrically connected pacemaker cells, adjacent interstitial cells and smooth muscle cells, and this unit thereby possesses sizeable electric capacity.

**Fig. S6.**
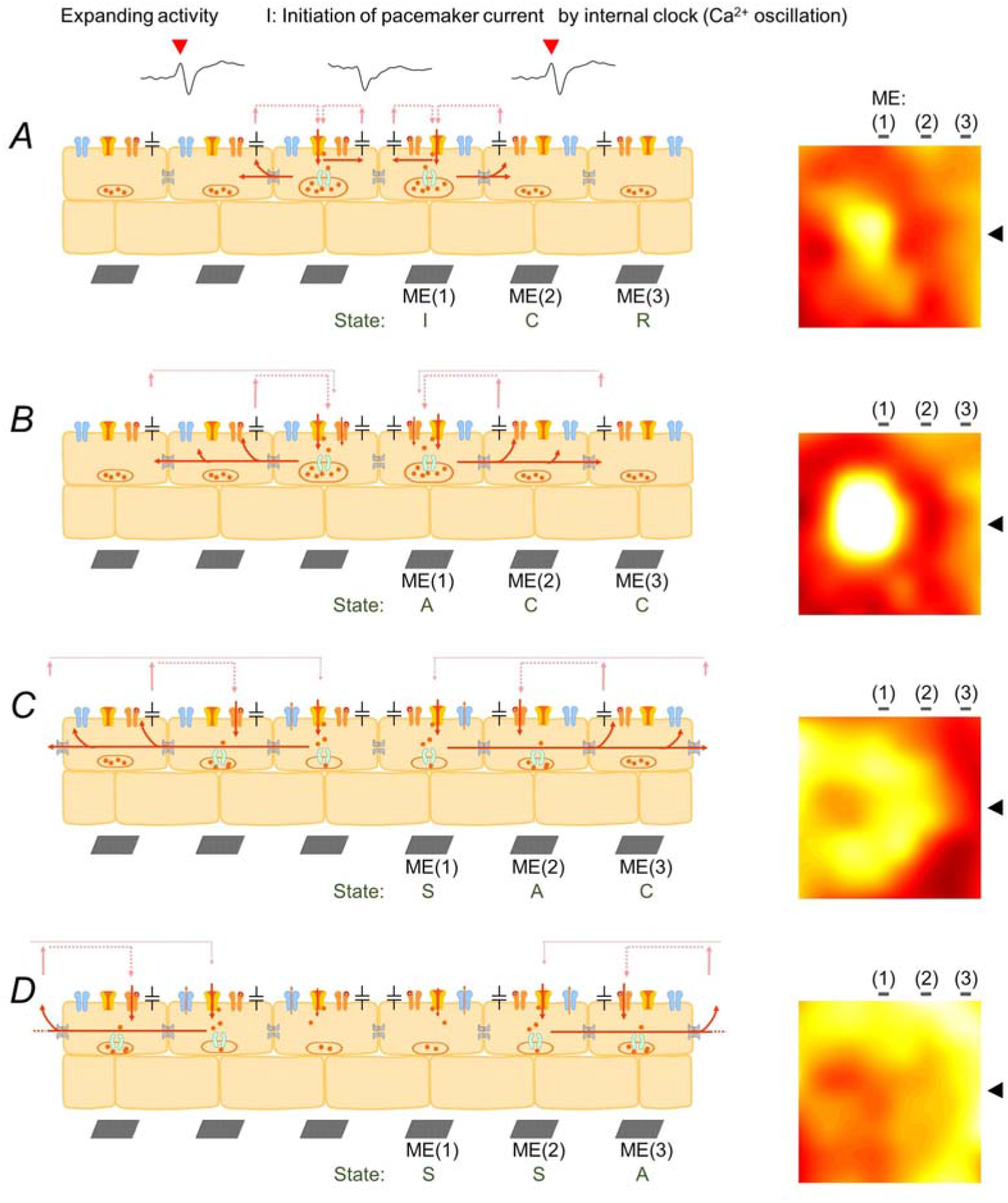
Schematic illustration showing the process underlying ‘expanding’ activity. (**A**) The cells in an initiating (‘I’) state near ME(1) employ the internal clock mechanism, such as intracellular Ca^2+^ oscillations owing to Ca^2+^ release channels in the endoplasmic reticulum, to invoke Ca^2+^-activated transmembrane pacemaker current (i.e., Cl^-^ channel inward current). Intercellular pacemaker current propagates toward the cells near ME(2) and charges their plasma membranes. The cells near ME(2) are in a charging (‘C’) state, while those near ME(3) are in a resting (‘R’) state. (**B**) Next, in the cells near ME(1), voltage-gated pacemaker current (a part of pacemaker current) is also activated in the cells near ME(1) so that these cells enter an activated (‘A’) state. Intercellular pacemaker current further propagates toward adjacent cells. The cells near ME(3) turn to the ‘C’ state. (**C**) In cells near ME(2), intercellular pacemaker current depolarizes the plasma membrane to the threshold and activates the voltage-gated inward current (a part of pacemaker current) so that these cells enter an activated (‘A’) state. On the other hand, the cells near ME(1) enter a slow oscillation (‘S’) state in which transmembrane outward K^+^ current and Ca^2+^-activated inward current (and/or voltage-gated inward current) are both activated. (**D**) The cells near ME(3) enter the ‘A’ state in which voltage-gated inward current is activated by intercellular pacemaker current, while the cells near ME(2) enter the ‘S’ state in which both outward and inward current are activated. Note that pacemaker potentials (negative transient potentials) near the initiating site lack a preceding positive potential (red arrowhead) which is prominent in field potentials observed in adjacent microregions. Black arrowheads indicate the position of MEs in the field potential image.

**Fig. S7.**
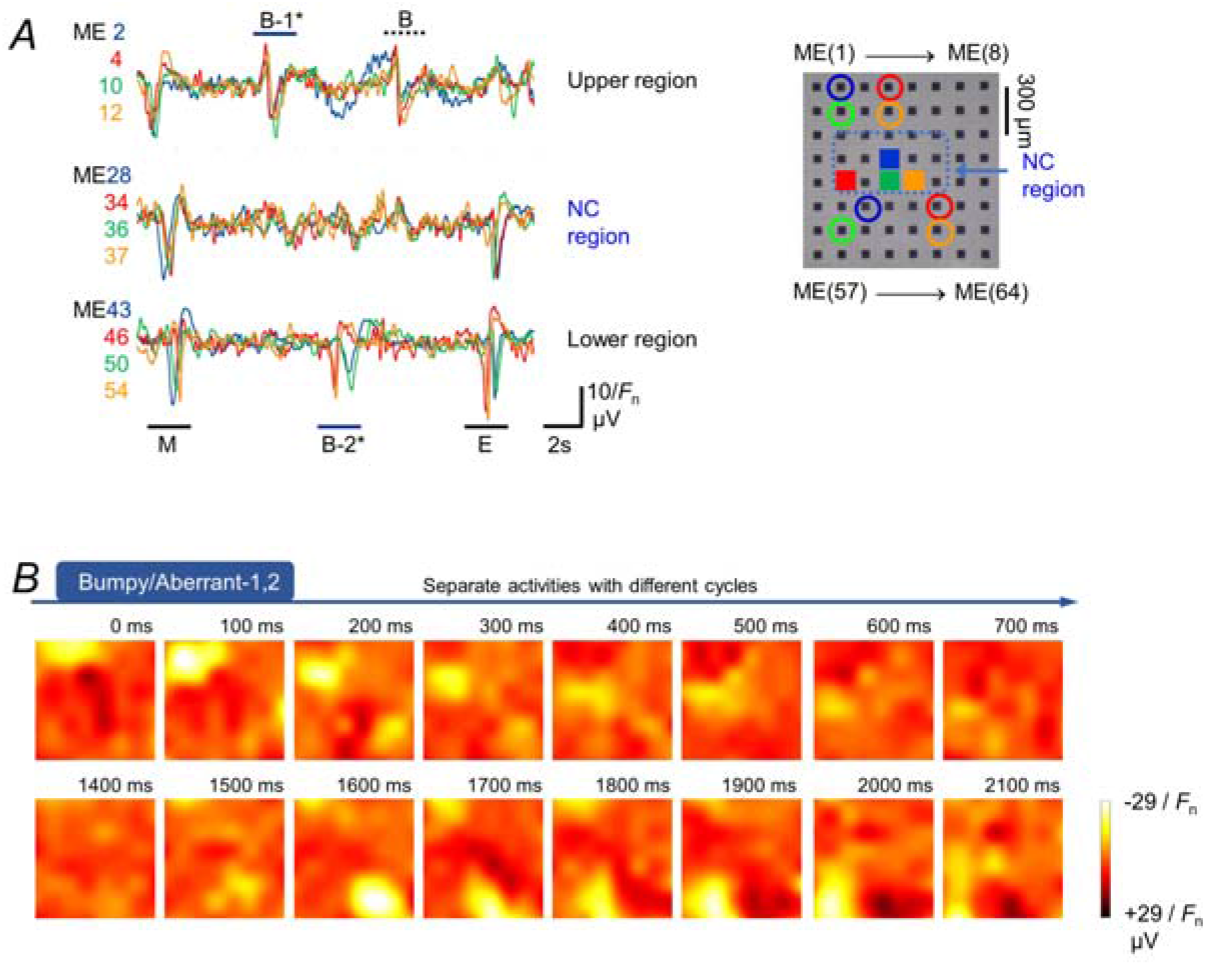
Pacemaker activities separated by a transient ‘non-conducting (NC)’ region, in a muscle sample showing an interconversion of pacemaker event patterns. (**A**) Field potentials recorded from different regions of the MEA. (**B**) Field potential images showing the two ‘bumpy/aberrant’ activities (B-1 and B-2) indicated in the field potential recorded in A. The B-1 and B-2 ‘bumpy/aberrant’ events occurred with different cycles and propagated locally in different regions (upper and lower, respectively).

**Fig. S8.**
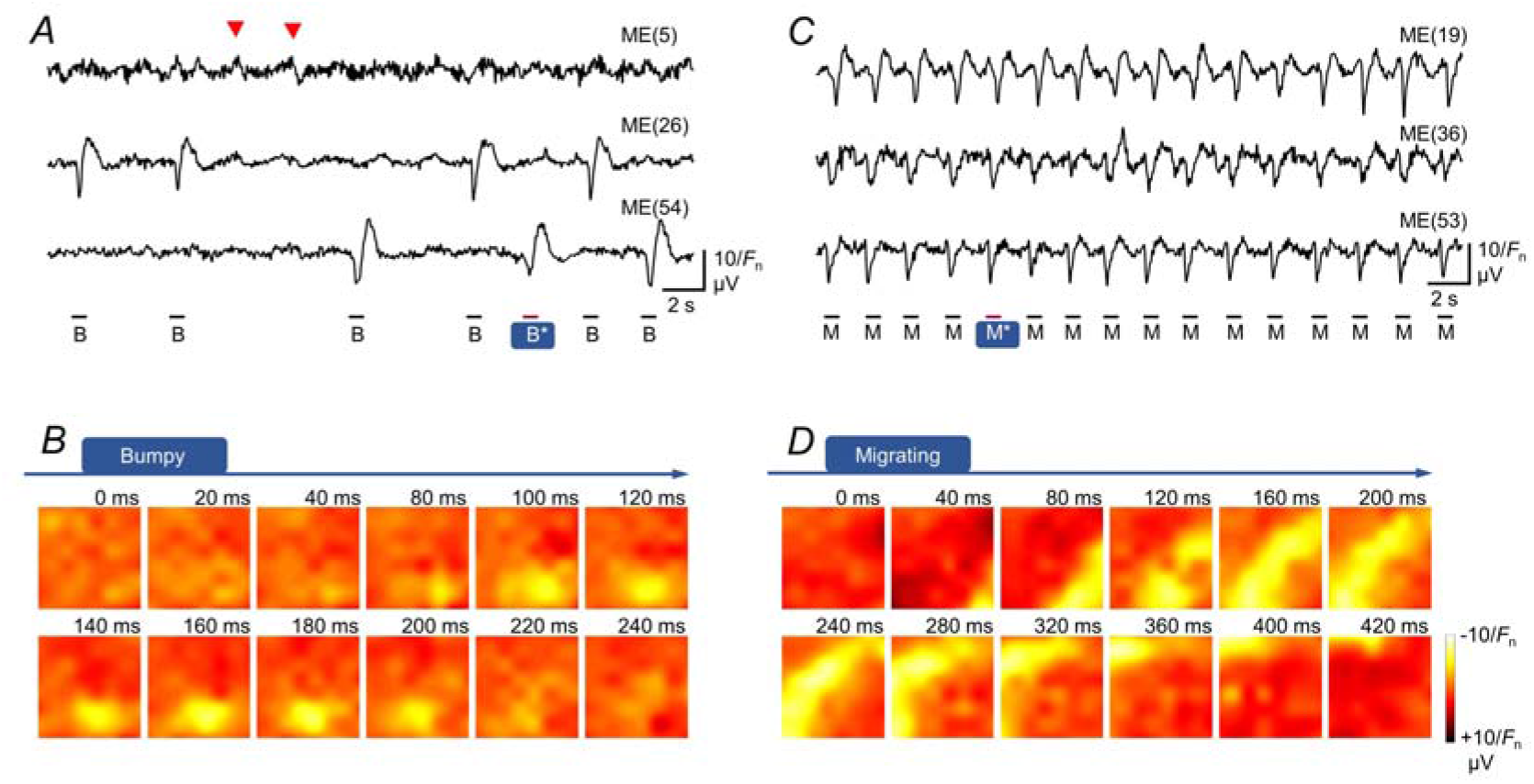
Modulation of micro-coordination patterns by 5-HT in a sample showing frequent ‘bumpy’ activity. Field potential traces recorded from different microelectrodes and corresponding field potential images obtained under control conditions (**A, B**) and in the presence of 5-HT (**C, D**) (an example of muscle sample showing frequent ‘bumpy/aberrant’ events in the control out of *N* = 9 experiments in nine samples). The field potential images correspond to the B* (‘bumpy/aberrant’ in the control) and M* (‘migrating’ in 5-HT) events indicated in the field potential traces. During the first half of the control recording period, premature activity-like potentials (red arrowheads) occurred between the ‘bumpy/aberrant’ events. The size of the potential image is 1050 × 1050 μm^2^.

**Fig. S9.**
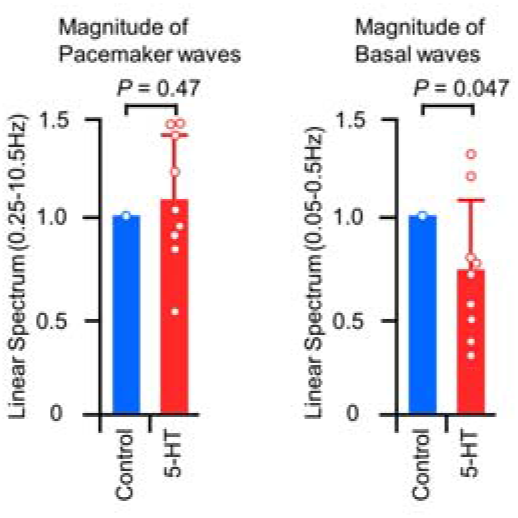
Effects of 5-HT on the magnitude of field potential waves. (**A**) Graphs showing the effect of 5-HT on the magnitude of the linear spectrum in the frequency range of pacemaker potential imaging (left: LS _0.25-10.5 Hz_, 1.08 ± 0.33 in the control, *P* = 0.47, *N* = 9) and in the frequency range of sub-basal waves (right: LS _0.05-0.5 Hz_, 0.73 ± 0.12 in the control, *P* = 0.047, *N* = 9).

## Supplemental Videos

The size of all videos is 1050 × 1050 μm^2^. The speed of videos was slowed four times.

**Supplemental Video SV1. Field potential video showing the ‘expanding’ pattern of a pacemaker event, corresponding to the data in Fig. 1F**.

SV1: Movie S1.mp4

**Supplemental Video SV2. Field potential video showing the ‘migrating’ pattern of a pacemaker event, corresponding to the data in Fig. 1F**.

SV2: Movie S2.mp4

**Supplemental Video SV3. Field potential video showing the ‘bumpy/aberrant’ pattern of a pacemaker event, corresponding to the data in Fig. 1F**.

SV3: Movie S3.mp4

**Supplemental Video SV4. Field potential video showing the ‘colliding’ pattern of a pacemaker event, corresponding to the data in Fig. 1F**.

SV4: Movie S4.mp4

**Supplemental Video SV5. Field potential video corresponding to Fig. 2A**. Three ‘expanding’ activities were initiated from distinct regions.

SV5: Movie S5.mp4

**Supplemental Video SV6. Field potential video showing two consecutive ‘migrating’ activities**. The ‘migrating’ activities propagated diagonally toward the lower-right region and longitudinally toward the upper region.

SV6: Movie S6.mp4

**Supplemental Video SV7. Field potential video showing consecutive pacemaker events in the control, corresponding to Fig. 4A**. A variety of micro-coordination patterns, including those of the ‘expanding’ pattern, occurred in the control.

SV7: Movie S7.mp4

**Supplemental Video SV8. Field potential video showing consecutive pacemaker events in the presence of 5-HT, corresponding to Fig. 4C**. ‘Migrating’ activities alone propagated diagonally toward the lower-left region, accompanied by an increase in the frequency.

SV8: Movie S8.mp4

**Summary Figure highlighting ‘expanding’ activity**

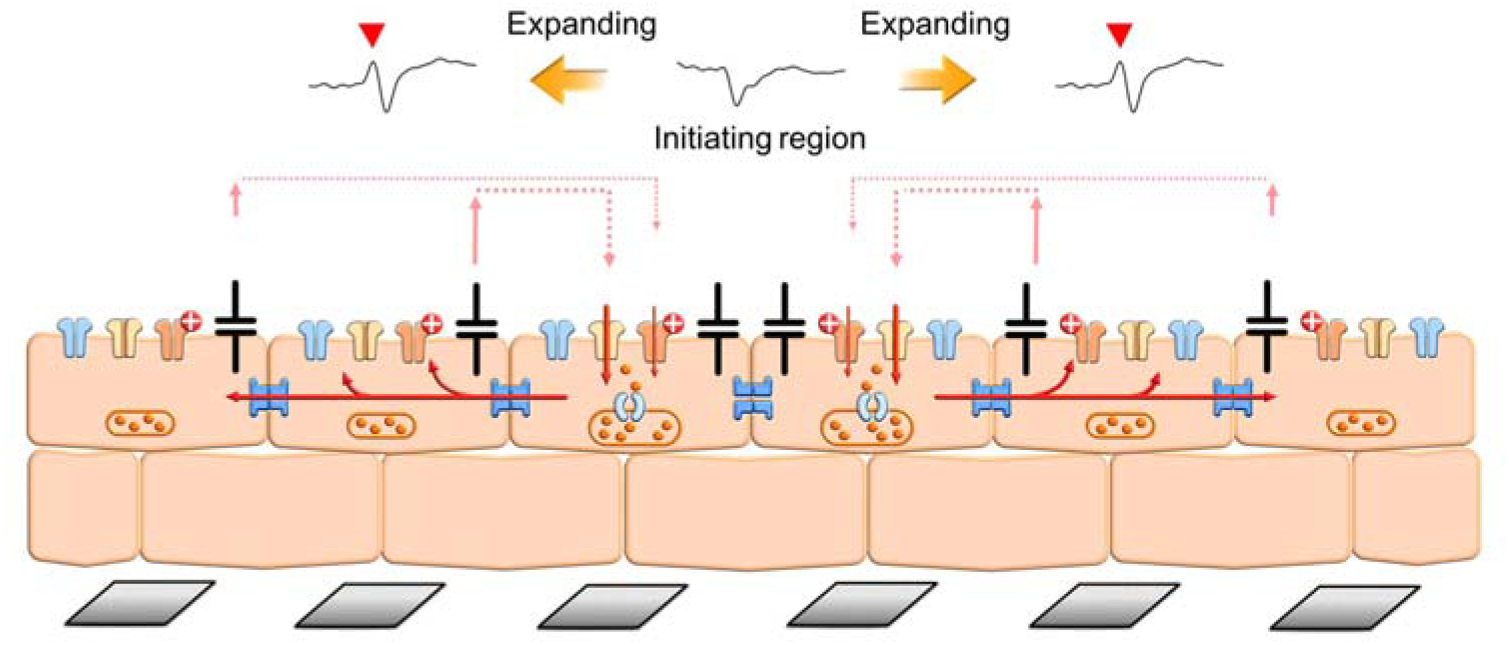

## Notes

### Competing Interest Statement

The authors have declared no competing interest.

### Summary of Updates

This version of manuscript is totally rewritten to emphasize the importance of methodological aspect, and is going to be submitted to Biosensors and Bioelectronics: X, as a revised manuscript.

